# Interpretable sequence-based machine learning consolidates candidate H3N2 hemagglutinin antigenic sites

**DOI:** 10.64898/2026.04.28.721429

**Authors:** Austin G. Meyer, Mauricio Santillana

## Abstract

Vaccine strain selection for seasonal influenza A(H3N2) depends on knowing which hemagglutinin (HA) substitutions are most likely to erode neutralizing antibody recognition, yet published antigenic site sets disagree substantially on which positions matter most. We applied interpretable gradient-boosted tree models with SHAP-based site attribution to two complementary hemagglutination inhibition (HI) datasets to produce a more consolidated ranking of candidate antigenic positions. Models trained on a Neher/Bedford benchmark dataset recover the canonical cluster-transition sites established by prior analyses. Moreover, after filtering the WIC dataset for confounding factors, our models recover the majority of positions from four major prior reference sets (Koel, Neher/Bedford, Harvey, and Shah) and improve concordance between rankings derived from the Neher/Bedford and WIC datasets. Rankings from our models also agree more strongly with models trained to predict sampling time or passage identity than with standard evolutionary metrics used to detect diversifying selection. Our results show that interpretable sequence-based models can provide a more integrative ranking of candidate antigenic positions across different data sources and modeling approaches. This work should aid efforts to prioritize H3N2 substitutions for epidemic surveillance.

**Significance Statement:** Every year, health authorities must update the seasonal flu vaccine to account for mutations in influenza A(H3N2) that allow the virus to escape existing immunity. Knowing which specific positions in the hemagglutinin protein drive this immune escape is essential for evaluating newly emerging variants, but published studies disagree substantially on which positions matter most. We show that interpretable machine learning models applied to two hemagglutination inhibition datasets, the Neher/Bedford benchmark dataset and the larger WHO Collaborating Centre dataset, can help to resolve the disagreements. The models recover canonical cluster-transition sites from the Neher/Bedford benchmark data, and show that our analysis approach with the WIC data improves concordance across several prior rankings produced from distinct datasets and modeling approaches. The resulting rankings provide a practical, consolidated reference for prioritizing hemagglutinin mutations most likely to affect vaccine effectiveness.

## 1 Introduction

Vaccine strain selection for seasonal influenza A(H3N2) depends on correctly prioritizing which amino acid substitutions in hemagglutinin (HA) are most likely to alter neutralizing antibody recognition. Since its emergence in 1968, H3N2 has continuously accumulated substitutions primarily in the HA head domain, the primary target of neutralizing antibodies. The substitutions erode cross-reactivity with vaccine-elicited antibodies and drive the need for frequent vaccine reformulation [1–3]. Global surveillance and vaccine-strain selection therefore rely heavily on hemagglutination inhibition (HI) assays that directly quantify antibody cross-reactivity between the vaccine strain and circulating virus strains [4, 5]. The utility of these assays is dependent on knowing which HA positions matter most. When a newly observed substitution occurs at a site with established antigenic significance, the appropriate response is clear: it deserves immediate serological follow up. When a substitution occurs at a position of uncertain importance, the appropriate response is more difficult. A more complete and reliable map of which HA positions are most strongly associated with H3N2 antigenic change would directly improve this decision process.

Despite substantial effort, such a map remains elusive, and published sets of important sites disagree more than one might expect. Early crystallographic and antibody-mapping studies defined five canonical H3 antigenic regions [6–9]. Cluster-transition analyses then narrowed the important sites to a small number of receptor-binding-proximal positions that explain major antigenic shifts [10, 11]. Subsequent additive titer models within phylogenetic frameworks extended the important set to approximately 15 positions [12, 13], a Bayesian structural-genomic analysis of the WHO Collaborating Centre (WIC) HI data prioritized positions using posterior inclusion probabilities [14], and a seasonal AdaBoost model trained on partially overlapping WIC data identified 30 positions through feature importance aggregated across influenza seasons [15]. These four reference sets (Koel, Neher/Bedford, Harvey, and Shah) overlap at their core but diverge considerably at other positions. That divergence likely reflects differences in dataset composition, time period, analytical method, and the degree to which laboratory passage artifacts contaminate the underlying HI measurements [16–20].

A separate class of approaches uses evolutionary variability as a proxy for antigenic importance. For example, sitewise dN/dS, the ratio of amino acid-changing to silent mutation rates, and Shannon entropy identify rapidly evolving or unusually variable positions [21–24], but neither directly measures antigenic effect. Moreover, both struggle with well-documented confounding from passage artifacts [17], structural constraints that limit site variability independent of antigenic pressure [25, 26], and rate estimation artifacts at short divergence times [27, 28]. Statistical and machine learning approaches offer a more direct route: sequence-based models have been applied to predict antigenic phenotype [29] and influenza fitness [30, 31], and interpretable methods have been used to prioritize biologically meaningful residues from protein-sequence data [32–34]. However, no study has systematically tested whether interpretable models can reduce disagreement among existing H3N2 reference site sets by applying consistent modeling assumptions, importance definitions, and confounder control.

Here, we address this gap directly. We apply interpretable gradient-boosted tree models (LightGBM) with SHAP-based importance attribution to two complementary HI datasets: the Neher/Bedford H3N2 dataset, which provides a clean benchmark against canonical cluster transition analyses, and the larger WIC H3N2 dataset, which covers more contemporary H3N2 evolution during the most recent decade. Without imposing an explicit evolutionary model, we filter the sequence data to reduce passage-artifacts and control for key confounders within the modeling framework. We then compare the resulting SHAP-based site rankings against all four reference sets, against independent rankings derived from sequence-only passage-history and collection-date prediction [35], and also against standard evolutionary metrics. We find that LightGBM+SHAP recovers the majority of the core positions identified by prior analyses while helping to reconcile several disagreements among them. The result is a more broadly concordant site ranking of candidate antigenic positions across benchmarks, datasets, and independent analytical frameworks.

## 2 Results

### 2.1 Interpretable machine learning HI-predictive models recover the canonical H3N2 cluster-transition sites

Our analysis focuses on two complementary LightGBM models that ask related but distinct biological questions. The site-state model asks whether the amino-acid states present at a given HA position in the virus–serum pair affect the predicted titer; it encodes each position with a binary change feature, a one-hot encoded virus amino acid (a separate binary indicator for each possible amino acid), and a one-hot encoded serum amino acid. The substitution model asks whether a specific directed amino acid change from the reference to the virus matters; it encodes each observed directed change at each position as a binary indicator. A temporal distance covariate was included in both encodings. Grouped cross-validation assigned all observations for a given virus to the same fold, preventing the same virus strain from appearing in both training and test data. See the Methods for further details.

The Neher/Bedford H3N2 dataset comprises 9,024 HI measurements retained after sequence matching, alignment, and quality control. The data cover major H3N2 antigenic cluster transitions from the late 1980s through the early 2010s (Table 1). For this dataset, the LightGBM models were trained to predict the residual standardized log_2_ HI titer after removing per-serum and per-virus additive offsets from full mature-HA sequence features. The site-state model achieved a mean cross-validation Spearman correlation of 0.708 and RMSE of 2.059 log_2_ titer units; the substitution model performed comparably (Spearman 0.695, RMSE 2.128; Table 1; Fig. S1). Both substantially outperformed a baseline using only temporal distance without sequence features (Spearman 0.594). Cross-validation performance was consistent across folds (Fig. S2).

**Table 1:** Predictive performance of the primary models. HI observations are virus–serum pairs retained for model fitting after matching both components to HA sequences and applying alignment quality control (see Methods). For the WIC dataset, 104,972 of 202,605 raw observations had matched HA sequences for both components, and 104,605 remained after HA1 alignment quality control. Performance values are mean grouped cross-validation RMSE and Spearman correlation between predicted and observed log_2_ HI titers across held-<u>out folds.</u>

| Dataset | HI observations | Model | RMSE | Spearman |
| --- | --- | --- | --- | --- |
| Neher/Bedford H3N2 | 9,024 | site-state | 2.059 | 0.708 |
|  |  | substitution | 2.128 | 0.695 |
| WIC H3N2 | 104,605 | site-state + context | 1.214 | 0.822 |
|  |  | substitution + context | 1.250 | 0.811 |
| WIC H3N2 (filtered) | 28,448 | site-state + context | 0.975 | 0.790 |
|  |  | substitution + context | 0.997 | 0.780 |

*Note:* The filtered WIC analysis excludes observations whose normalized virus or reference passage class is EGG, MIXED, or UNKNOWN. Because filtering changes the response distribution, RMSE and Spearman values are not directly comparable between the unfiltered and filtered rows (see Methods). “Context” indicates that assay-context covariates are included alongside sequence features after an additive avidity/potency correction (see Methods). Sequence numbering uses mature HA numbering, excluding the 16-residue signal peptide. Full results including patristic-distance sensitivity analyses are in Table S1.

SHAP (SHapley Additive exPlanations) values quantify how much each sequence feature contributes to the model’s predicted titer for a given virus–serum pair; sites with large mean absolute SHAP values are those whose sequence states most consistently predict antigenic distance. A site’s mean signed SHAP further indicates the direction of that contribution: positive values mean that sequence differences at that site push the prediction toward greater antigenic distance (plausibly from immune escape), while negative values mean the site’s state is associated with preserved cross-reactivity. Because HA substitutions are correlated along clades, these site rankings should be interpreted as model-based attribution of predictive signal rather than proof that each site independently causes antigenic change.

To provide context for evaluating the overlaps reported below, the four reference sets include Koel (7 positions), Neher/Bedford (15 positions), Harvey (15 positions), and Shah (30 positions). The Harvey and Shah sets both derive from the WIC HI data, but share only 8 of their combined positions. Under a hypergeometric null, two independently drawn sets of 30 from 328 HA1 sites would overlap by approximately 2.7 sites on average. Therefore, the overlaps should be interpreted against this baseline.

SHAP values from the site-state model prioritized sites in the canonical head-region positions associated with H3N2 cluster transitions (Fig. 1). The top-ranked positions, including 189, 135, 145, 121, 193, and 156, cluster in and around antigenic sites A and B, directly overlapping the receptor-binding-proximal region where cluster-defining substitutions accumulate [7, 11]. Sites 159, 62, and 186 also ranked highly; each is found in the Neher/Bedford analysis and at least one of the prior WIC analyses. Of the seven Koel cluster-transition positions (145, 155, 156, 158, 159, 189, 193) [11], our models recovered six among the top 30 site-state model sites, and all seven in the top 30 by the substitution model (Table S2). Of the 15 Neher/Bedford reference positions, 11 appeared in our top 30; 11 of 15 Harvey positions and 12 of 30 Shah positions likewise appeared. All of these overlaps substantially exceed chance expectation. Several of the leading sites (e.g., 189, 135, 145, 193) carry positive mean signed SHAP, suggesting that sequence differences at these positions are consistently associated with increased antigenic distance. By contrast, sites 133 and 278 rank highly by absolute SHAP but carry negative mean signed SHAP, indicating that the amino acid states at these positions are informative in the direction of preserved cross-reactivity rather than antigenic escape. A fold-wise stability analysis confirmed that the leading positions were recovered consistently across cross-validation folds, with the top 15 sites showing particularly narrow rank distributions (Fig. S3). A secondary grouped-virus holdout evaluation, in which 10% of virus strains were withheld together, confirmed that the sequence models generalize to previously unseen test viruses rather than memorizing training measurements (Fig. S4).

**Figure 1:**
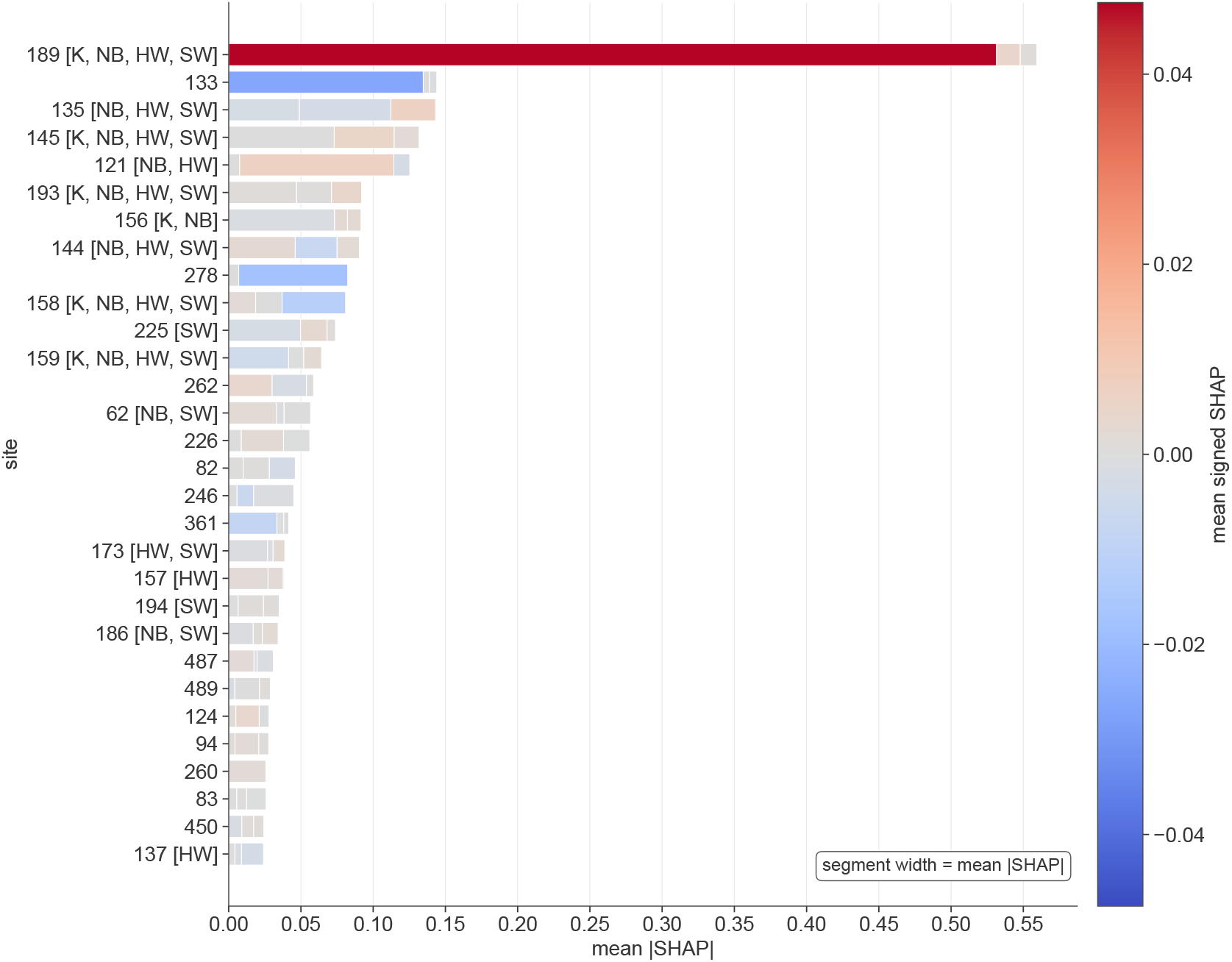
Site-level SHAP decomposition for the site-state model trained on the Neher/Bedford dataset. Bars show the top 30 ranked mature-HA sites by mean absolute SHAP value across grouped-cross-validation folds. Bar segments indicate the contribution of site-change, virus-state, and serum-state components to the aggregated site SHAP score. Brackets at right annotate each site with membership in the Koel reference set (K; 7 positions), the Neher/Bedford reference set (NB; 15 positions), the Harvey/WIC reference set (HW; 15 positions), and the Shah/WIC reference set (SW; 30 positions), with coloring matched to the respective literature source. The leading positions include canonical cluster-transition sites such as 189, 145, 156, 158, 159, and 193. All site numbers use mature HA numbering, excluding the N-terminal 16-residue signal peptide.

As a sensitivity test, adding patristic distance (the total branch length separating two sequences on a phylogenetic tree) as an additional covariate did not meaningfully change either predictive performance (Spearman 0.708, RMSE 2.054 vs. Spearman 0.708, RMSE 2.059 without patristic; Table S1) or the top-ranked site set (full patristic sensitivity reported in the Sensitivity Analyses section below). The near-identical Spearman values and the marginal RMSE reduction indicate that patristic distance absorbs little variance beyond what sequence features already capture, and in general the site importances are unchanged. The site-state model trained on the Neher/Bedford dataset thus provides a clean benchmark reconstruction: it recovers the canonical H3 cluster-transition site set without requiring structural data, fitness models, or phylogenetic correction.

### 2.2 Passage filtering and interpretable machine learning clarify the biological signal in the WIC dataset

The WIC H3N2 dataset spans matched virus–serum HI observations collected over multiple decades, with most modeled observations concentrated after 2010. The WIC data cover a much broader range of laboratories, reference panels, and virus passage contexts than the Neher/Bedford collection. The raw dataset contains 202,605 HI observations; 104,972 had matched HA1 sequences for both virus and serum components, and 104,605 observations (52%) remained after alignment quality control for modeling. Egg passage of H3N2 routinely introduces amino acid substitutions at receptor-binding-proximal positions such as 194, 219, 246, and 248 that alter HI titer patterns independently of the biologically circulating virus [16, 18–20]. When egg-passaged viruses and reference antisera are included in our models, these passage-specific substitutions appear as predictive features but are mechanistically unrelated to the antigenic phenotype of the circulating strain.

Applying a passage-history filter, which excludes observations whose normalized virus or reference passage class is `EGG, MIXED`, or `UNKNOWN`, retained 28,521 HI measurements after sequence matching; 28,448 of these had aligned HA1 sequences available for feature construction and entered the filtered modeling analysis. This filtering trades representativeness for interpretability: the retained subset is smaller and enriched for more recent, cell-associated observations, but it removes the passage classes most likely to confound residue-level attribution (Table S3). The site-state model trained on the passage-filtered WIC data achieved a mean cross-validation Spearman correlation of 0.790 and RMSE of 0.975 (Table 1; Fig. S5). The corresponding site-state model trained on the unfiltered WIC dataset shows a nominally higher Spearman correlation (0.822) and a higher RMSE (1.214). The two metrics are not directly comparable across the filtered and unfiltered datasets because filtering changes the response distribution. The lower filtered Spearman reflects a narrower range of titer variation after removing passage-heterogeneous observations rather than worse model fit. The lower filtered RMSE reflects the removal of passage-induced noise rather than better per-observation accuracy. Given the tighter predicted-versus-actual scatter (Fig. S6) and cross-study concordance analysis below, the models trained on the passage-filtered WIC data are our preferred primary models because they remove known confounding by passaging and provide the clearest biologically credible site ranking.

The site-state model trained on the passage-filtered WIC data identified positions 144, 225, 159, 140, and 145 as the leading sites associated with antigenic variation, with sites 62, 158, and 189 also among the top 15 (Fig. 2). The prominence of site 144 is consistent with prior analyses identifying residue 144 and the surrounding site-A loop as antigenically important in contemporary H3N2 evolution [14]. Site 225 ranked second overall despite falling outside the classical head-domain epitope designations; we discuss the possible biological significance in the Discussion. Fold-wise stability confirmed consistent recovery of important sites across folds (Fig. S7). A diagnostic check confirmed that the additive context correction successfully removed the bulk of assay-specific titer variation (laboratory, passage class, and antiserum differences) before sequence features were interpreted (Fig. S8). A secondary grouped-virus holdout confirmed that sequence models outperformed the context-only baseline (without sequence data) (Fig. S9).

**Figure 2:**
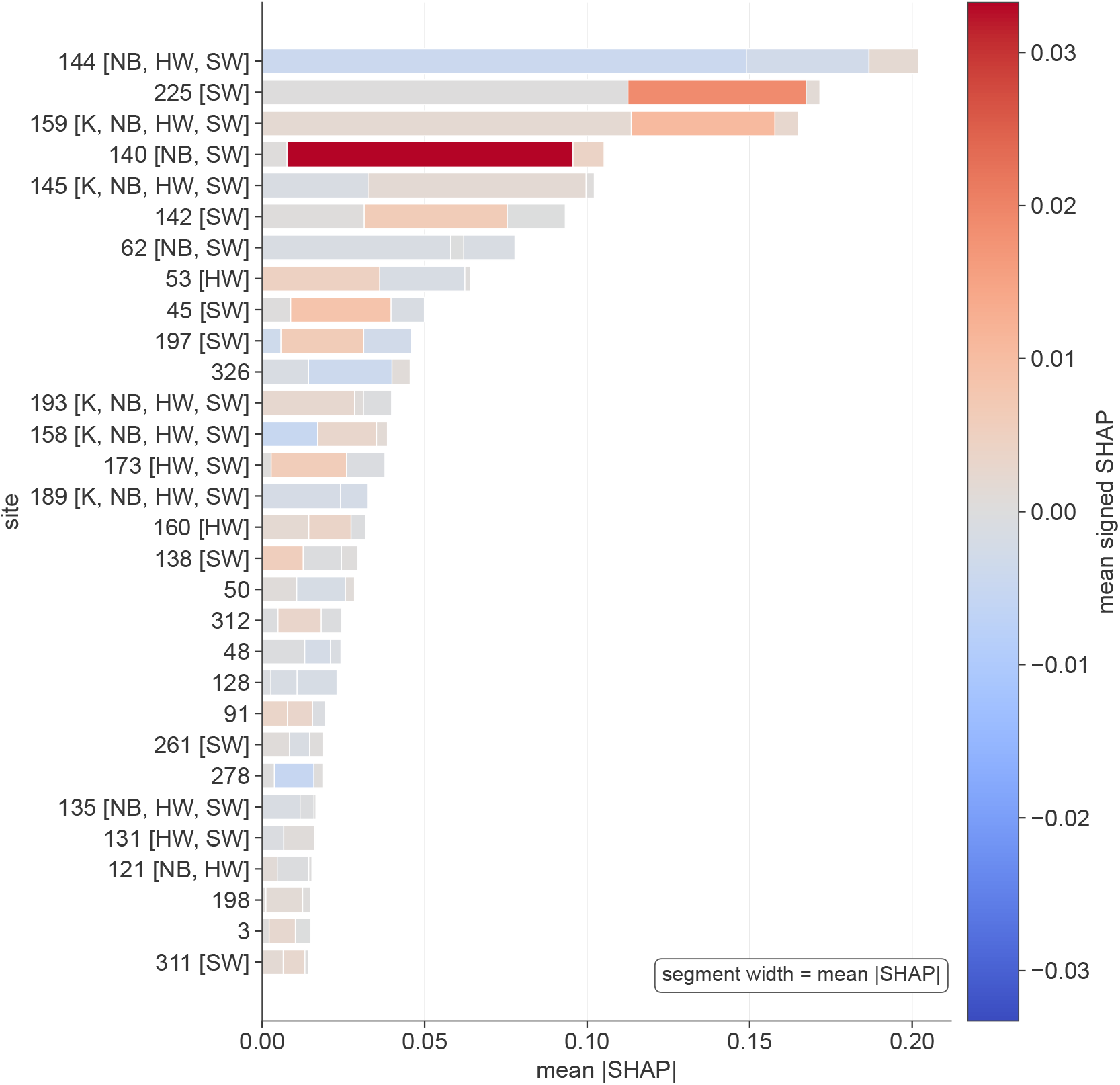
Site-level SHAP decomposition for the site-state model trained on the passage-filtered WIC data. Bars show the top 30 ranked HA1 sites by mean absolute SHAP value across grouped-cross-validation folds. Bar segments and bracket annotations follow the same conventions as Fig. 1. The filtered analysis excludes rows whose normalized virus or reference passage class is `EGG, MIXED`, or `UNKNOWN`. The leading sites (144, 225, 159, 140, 145, 62, 158, and 189) reflect the canonical head-region antigenic signal and the post-2010 H3N2 substitutions that drove clade transitions during the period covered by the WIC dataset; site 225 ranks second overall but falls outside the classical head-domain epitope designations.

Models trained on the unfiltered WIC dataset returned noisier site rankings. The two-cluster structure visible in the unfiltered predicted-versus-actual scatter (Fig. S10) reflects the coexistence of multiple passage contexts, especially egg-associated and cell-associated observations, occupying distinct regions of titer space; passage-associated positions ranked more prominently alongside canonical antigenic sites in the unfiltered decomposition (Fig. S11). Adding patristic distance to the site-state model trained on the passage-filtered WIC data provided marginal improvement (Spearman 0.795 vs. 0.790, RMSE 0.964 vs. 0.975; Table S1), indicating that interpretability depends more on the data quality of the input observations than on phylogenetic correction.

### 2.3 Cross-study synthesis identifies a convergent set of antigenically informative H3N2 sites

The four prior reference sets, Koel, Neher/Bedford, Harvey, and Shah, agree at their core but show limited pairwise overlap elsewhere. As noted above, Harvey and Shah share only 8 of their combined positions despite drawing on similar WIC HI data; Koel and Neher/Bedford share 6 or 7 Koel positions, but pairwise overlap between more distant pairs is considerably lower. To determine how well our models can bridge this disagreement, we compared the top 30 sites from each primary model against all four benchmark sets and against several external sequence-only rankings that do not use HI data (Fig. 3; benchmark overlap counts in Table S2).

**Figure 3:**
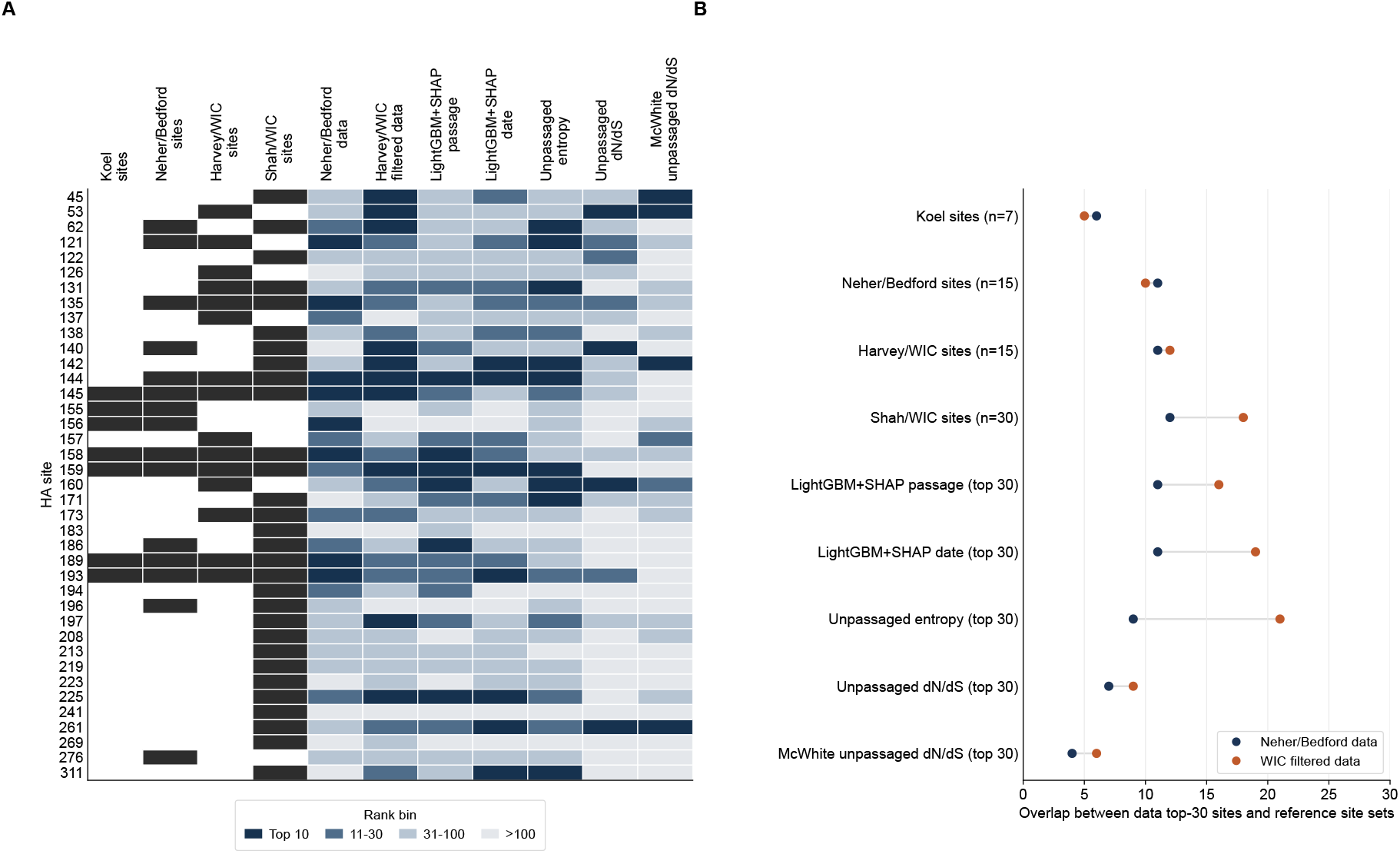
Cross-study concordance showing how the SHAP-based models relate to prior antigenic site sets. **(A)** Site-by-study comparison across the union of Neher/Bedford and Harvey reference positions. Each row is a site; columns indicate rank tier for the site-state model trained on the Neher/Bedford data, the site-state model trained on the passage-filtered WIC data (WIC_filt_), and each external ranking. Darker shading indicates a higher rank within the top 30; membership in the Koel, Neher/Bedford, Harvey/WIC, and Shah/WIC reference sets is shown in the rightmost columns. **(B)** Overlap of the top-30 site-state site sets from models trained on the Neher/Bedford and passage-filtered WIC data with each benchmark or external ranking. Point distance from the left indicates the number of matching sites out of the reference set size (7 for Koel; 15 for Neher/Bedford and Harvey/WIC; 30 for Shah/WIC and sequence-based external rankings). The distance between points on the same row indicates the relative concordance of models trained on the two distinct datasets. The site-state models trained on the Neher/Bedford and passage-filtered WIC data capture many core positions from each prior analysis and largely recover the same sites across input datasets. Harvey and Shah, which analyzed related WIC-derived data, overlap at 8 sites; the model trained on passage-filtered WIC data overlaps with 12 Harvey sites and 18 Shah sites. The model trained on passage-filtered WIC data also overlaps with 10 Neher/Bedford reference sites, compared with 8 for the Shah set and 8 for the Harvey set.

The two site-state analyses showed complementary strengths. The model trained on the Neher/Bedford data recovered the canonical cluster-transition sites most completely, capturing nearly all Koel and Neher/Bedford reference positions in its top 30 (Table S2). The model trained on the passage-filtered WIC data provided the broadest overall agreement; it matched a slightly larger fraction of the Harvey set and showed greater agreement with the Shah seasonal-model positions (Table S2) than the model trained on the Neher/Bedford data. Importantly, site-state models trained on either dataset overlap with each of the four prior reference sets at rates that, in most cases, exceed the pairwise overlap those reference sets show with each other. This overlap suggests that our approach captures a broader consensus of antigenic sites across datasets and methods.

The SHAP-based rankings showed strong agreement with rankings derived independently from sequence data alone, without HI measurements. The top 30 sites from the site-state model trained on the passage-filtered WIC data overlapped substantially with sequence-only LightGBM+SHAP site rankings derived from models that predict passage history and collection date [35], and with the most variable positions in unpassaged H3N2 sequences as measured by Shannon entropy (Fig. 3B). This convergence across HI-based and sequence-only approaches supports the interpretation that the prioritized sites reflect a genuine biological signal rather than an artifact of any single dataset or modeling strategy. By contrast, overlap with sitewise dN/dS was much weaker, and overlap with the McWhite et al. unpassaged dN/dS ranking [17] was the weakest of all comparisons; we discuss this pattern further in the Discussion.

### 2.4 Sensitivity analyses support the primary interpretation

Two additional sensitivity analyses confirmed the robustness of the main conclusions. Including the full, unfiltered WIC observations yielded a noisier and less biologically interpretable site ranking (Fig. S10, S11), consistent with the well-documented effect of passage heterogeneity on site-level attribution. Adding patristic distance as a covariate in both primary models produced RMSE improvements of roughly 1% or less and did not alter the leading ranked sites (Table S1; Fig. S12, S13, S14, S15). These minimal gains indicate that patristic distance explains little variance beyond what sequence features already capture and that the core site attributions are not artifacts of omitting a phylogenetic correction term.

## 3 Discussion

We show that interpretable gradient-boosted tree models with SHAP importance attribution help reconcile the partially overlapping site sets from four prominent H3N2 antigenic analyses

— Koel, Neher/Bedford, Harvey, and Shah — into a more integrative ranking. Our model trained on the Neher/Bedford data recovers virtually all of the canonical cluster-transition sites [11, 13], confirming that the framework faithfully reproduces the historical benchmark signal against which any new analytical approach should be tested. In addition, modeling the filtered WIC data yields a ranking with stronger overlap across all four reference sets than those sets show with each other [14, 15], and does so without requiring protein structural information or explicit phylogenetic modeling.

The practical advantage of LightGBM+SHAP over prior frameworks is flexibility. Additive titer models, including the Neher/Bedford framework, represent the genetic component of antigenic change as a sum of branch or substitution effects plus confounding terms [12, 13]. This works well when antigenic effects are approximately additive, but it does not naturally accommodate cases in which the effect of a substitution depends on genetic background, as is known to occur at positions 145 and 155 [36, 37]. Bayesian structural-genomic models such as Harvey et al. account for uncertainty more formally and incorporate phylogenetic and structural information [14], but they require substantially heavier computation and a more prespecified probabilistic framework. Gradient-boosted tree models, like LightGBM, learn flexible nonlinear mappings from paired virus–serum sequence features to HI outcome, while SHAP provides site-level attribution tied directly to the measured phenotype. The two sequence encodings used here provide reinforcing evidence: the substitution encoding shows slightly stronger recovery of the Koel set, whereas the site-state encoding is especially useful in the WIC analysis where contemporary variation occurs across diverse amino acid backgrounds at the same sites.

Our site rankings are most informative when viewed jointly across datasets and encodings. The models built on the filtered WIC data extend the historical cluster-transition signal toward contemporary variation, with high rank for sites 144 and 140. That is consistent with the greater prominence of the antigenic A region in the post-2010 evolutionary background sampled by WIC [14, 15]. Identification of site 62 in both primary site-state models indicates that the broader ranking retains the site-E signal from earlier cluster-transition literature [7, 13].

Site 225 deserves specific attention. It falls outside of the classical head-domain epitope regions, and it has not been found in prior computational antigenic rankings (though it was identified in the Shah set), yet it ranked second overall in our filtered WIC model. A plausible biological connection exists: Chambers et al. identified antigenic-site-B mu-tations as the principal drivers of the 2014–2015 H3N2 vaccine mismatch, but the 3C.2a and 3C.3a drift backgrounds they experimentally dissected also carried N225D relative to A/Texas/50/2012 [38]. Given that site 225 is proximal to the receptor-binding site, its high rank may reflect a contribution to avidity-linked antigenic phenotypes rather than a classical epitope effect. This would be a specific hypothesis that could be directly tested through experimental follow up.

The agreement between SHAP-based rankings and several independent sequence-only rankings [35] strengthens the case that the prioritized sites reflect a genuine biological signal rather than artifacts of any single HI dataset or modeling strategy. At the same time, this convergence should not be overinterpreted as proof of antigenic causality: sites could rank highly in multiple analyses because of shared evolutionary confounding or correlated substitution patterns rather than because they directly alter antibody recognition. However, it does indicate that the signal is not dataset-specific. The weak overlap with sitewise dN/dS reinforces a related point: dN/dS identifies codons under diversifying selection but does not distinguish whether that selection is antigenically consequential [21, 23, 26], and it is further shaped by structural constraints that limit variation independently of antigenic pressure [25, 39]. By contrast, our SHAP-based importance ranking is tied directly to an experimentally measured HI outcome, making it better aligned with the surveillance task of prioritizing substitutions by likely antigenic effect.

Two practical modeling choices also emerged from our analysis. First, passage filtering matters. Applying the WIC passage filter, which removes rows whose normalized virus or reference passage class is egg, mixed, or unknown, produced a site ranking with improved concordance across benchmarks and provided a clearer biological interpretation. This is consistent with a large body of work showing that egg adaptation introduces substitutions at receptor-binding-proximal positions that can confound residue-level analyses [16–20]. Second, adding patristic distance as a covariate provided at most marginal improvement (about a 1% RMSE gain in the site-state model trained on the filtered WIC data, and less in the site-state model trained on the Neher/Bedford data) and did not alter the leading site rankings. This finding suggests that it may be possible to sufficiently account for phylogeny within a flexible nonlinear model without needing to explicitly specify an evolutionary framework.

Several limitations are important to note. Both datasets reflect the strain selection biases and assay preferences of the surveillance systems that generated them, and neither provides a complete sample of the H3N2 sequence–titer landscape. The filtered WIC analysis retains only about one quarter of the sequence-matched observations that pass alignment quality control (28,448 of 104,605), and some excluded measurements may carry biological signal that is lost. In addition, SHAP-based attribution identifies sites that most consistently predict measured HI titers but does not provide experimental proof of mechanism. Highly ranked sites could reflect correlated evolutionary processes, sampling biases, assay-specific effects, or SHAP credit distributed among correlated clade features rather than direct antibody-contact roles. The current feature encoding does not explicitly capture pairwise or structural interaction terms, which may matter at positions such as 156 and 158 [36, 37], nor does it represent glycosylation at N-linked sites which can add antigenic heterogeneity not captured by amino acid states alone [40–42]. Finally, the choice of aggregating absolute rather than signed SHAP values is one of several encoding decisions whose optimality remains an open question. Using signed values would, in our case, drop several sites with large negative contributions (including sites 82, 133, and 278), and likely improve concordance further. However, identifying the best aggregation scheme for this application warrants dedicated investigation.

For surveillance applications, positions that rank highly across both datasets, both encodings, and multiple reference comparisons represent the strongest candidates for antigenic significance and should be prioritized for serological follow up when observed in newly circulating variants. Positions prominent in one dataset but not the other may reflect temporal, dataset-specific variation, or be artifacts of specific modeling choices, and warrant experimental characterization before interpretation. When coupled with deep mutational scanning [43, 44] and antigenic cartography [3], the consolidated site rankings developed here provide a practical filter for directing experimental resources toward the substitutions most likely to affect the match between circulating H3N2 viruses and existing vaccine-elicited immunity [5, 13].

## 4 Materials and Methods

### 4.1 Data sources and sequence processing

The Neher/Bedford H3N2 HI dataset and matched HA sequences were obtained from the Neher/Bedford antigenic and phylogenetic analysis framework [13]. The WIC H3N2 HI dataset and passage annotations were assembled from the WHO Collaborating Centre surveillance data described in Harvey et al. [14].

For the Neher/Bedford dataset, HA sequences were retrieved from GISAID [45, 46] using isolate identifiers matched to the HI records, with NCBI GenBank as a fallback for isolates not resolvable through GISAID. The final Neher/Bedford modeling dataset retained 9,024 HI observations and 402 unique HA sequences after alignment quality control.

The WIC dataset required a multi-stage sequence search because the raw 202,605 HI observations reference 16,223 unique virus and reference serum strains, the majority of which are not indexed under consistent accession numbers in public databases. Sequences were searched through three sources applied in priority order: (1) the curated HA1 alignment distributed with the Harvey et al. analysis [14] (retrieved from the public code repository), which provided 1,738 matched sequences; (2) GISAID, queried by strain name [45, 46] and matched by exact and normalized strain-name lookup, which provided 2,501 additional sequences; and (3) NCBI GenBank and the BV-BRC (Bacterial and Viral Bioinformatics Resource Center) public databases, queried by strain name via the Entrez API and BV-BRC REST API, which yielded a small number of accession-linked manifest rows but no sequences in the final local FASTA used for modeling. The final modeling inputs therefore contained 4,239 resolved strains from the Harvey repository and GISAID, yielding 104,972 HI observations with sequences available for both the virus and reference serum components. The observation count is larger than the strain count because each resolved virus or reference serum strain can appear in many virus–serum pairings. After alignment quality control, 104,605 observations remained and were used for modeling. When multiple GISAID isolates corresponded to a given strain name, a canonical isolate was selected by preferring HA1-complete sequences (*≥* 328 residues), then lower-passage material (unpassaged *>* cell *>* egg), then lower sequence ambiguity, then the most common variant among HA1-complete candidates.

For the Neher/Bedford dataset, retrieved HA protein sequences were aligned to the A/HongKong/1/1968 reference sequence using MAFFT [47] in automatic mode, the N-terminal 16-residue signal peptide was removed, and the 550-residue mature HA sequence was retained for modeling. For the WIC dataset, HA1 sequences were extracted from the final protein FASTAs, aligned with MAFFT, and mapped to mature HA positions using the retained HA1 alignment. The two datasets were processed separately, but all reported positions use the same mature HA numbering convention, excluding the signal peptide. Sequences with more than 5% gap positions or more than 2% ambiguous residues were excluded. Full details of sequence retrieval, strain-name matching, GISAID canonical isolate selection, alignment, and numbering are provided in Supplementary Methods S1.1 and S1.2.

### 4.2 Titer preprocessing and avidity/potency correction

For both datasets, each titer was first standardized against the homologous titer of the corresponding serum (the titer measured against the virus used to raise the antiserum), so that larger values indicate greater cross-reactivity loss against heterologous viruses. Censored values reported as *<*X or *>*X were converted to X/2 and 2X before log_2_ transformation.

For the Neher/Bedford dataset, a simple additive correction using serum identity and virus strain as two categorical factors was applied to remove per-serum and per-virus titer

offsets before sequence features were interpreted. For the WIC dataset, passage history was used both to define a filtered subset and to account for laboratory-specific and assay-specific titer offsets. In the filtered analysis, observations were excluded when either the virus or reference serum had a normalized passage class exactly equal to EGG, MIXED, or UNKNOWN, retaining 28,521 observations after sequence matching; 28,448 remained after requiring aligned HA1 sequences for feature construction. An additive avidity/potency framework was then defined using six categorical factors fitted jointly by ordinary least squares: reference serum identity, assay-date label, virus passage class, reference serum passage class, antiserum class (ferret, human, or other), and red blood cell type. Full-data fitted values and residuals were saved as diagnostics, but during model training the additive layer was re-estimated within each split and models were trained to predict standardized titer relative to that split-specific additive baseline. The same confounding-factor framework was used for both the full and filtered WIC analyses. Full details of the residualization procedure and factor definitions are in Supplementary Methods S1.3.

The supervised learning target was therefore not the raw titer. For Neher/Bedford, sequence models predicted standardized titers after removal of per-serum and per-virus additive offsets. For WIC, the additive context layer was re-estimated within each training split, and the sequence models predicted standardized titer relative to that split-specific additive baseline. Reported held-out predictions and SHAP values therefore reflect sequence features plus allowed context covariates, with nuisance offsets estimated only from training data.

### 4.3 Feature encodings and model fitting

Two complementary sequence encodings were applied to each dataset. For Neher/Bedford, positions span the 550-residue mature HA sequence; for WIC, they span the 328-residue HA1 alignment. The site-state encoding represents each position with three feature groups: a binary change indicator (does the virus amino acid differ from the serum amino acid at this site?), a one-hot encoding of the virus amino acid state, and a one-hot encoding of the serum amino acid state. The substitution encoding represents each observed directed amino acid change from serum to virus at each position as a binary indicator. For models trained on the Neher/Bedford dataset, a temporal distance covariate (absolute calendar-year difference between virus and serum) was included. For models trained on the WIC dataset, the sequence features were augmented with temporal distance plus assay-context covariates describing virus passage class, reference passage class, whether each passage annotation was known, antiserum class, red blood cell type and species, and oseltamivir presence and concentration. Patristic distance was added as an additional covariate in the sensitivity variants (see below). Full encoding details are in Supplementary Methods S1.4.

Gradient-boosted tree models were trained using LightGBM [48] with up to 1,000 boosting iterations (learning rate 0.1, maximum tree depth unconstrained, minimum child samples 20, number of leaves 31, no regularization). Early stopping with a patience of 50 rounds was applied per cross-validation fold. Models were evaluated using five-fold grouped cross-validation in which all HI observations for a given virus strain were assigned to the same fold, preventing leakage from closely related viruses. A secondary grouped-virus holdout evaluation, implemented by withholding 10% of virus strains at once, was also performed (Figs. S4, S9). Validation procedures are described in Supplementary Methods S1.5.

### 4.4 Patristic distance computation

For the patristic-distance sensitivity analyses, maximum-likelihood phylogenies were estimated from the aligned mature-HA sequences for Neher/Bedford and from the aligned HA1 sequences for WIC using FastTree [49] under the WAG amino-acid substitution model with gamma-distributed rate variation (`-wag -gamma`). Pairwise patristic distances between each virus tip and the corresponding reference serum tip were extracted from the reconstructed Newick tree using Biopython’s `tree.distance()` function [50] and included as an additional covariate alongside sequence features. The rationale for treating these models as sensitivity analyses rather than primary analyses is described in Supplementary Methods S1.8.

### 4.5 SHAP attribution and site ranking

SHAP values [51] were computed for all site-state features using the TreeExplainer algorithm. For the Neher/Bedford dataset, values were aggregated over all held-out predictions in each cross-validation fold; for the larger WIC analyses, up to 2,500 held-out rows were sampled per fold before aggregation. For each site, the mean absolute SHAP values of the site-change, virus-state, and serum-state feature groups were summed to produce an aggregate per-site importance score over the pooled held-out observations used for SHAP, which defined the final ranking. Fold-wise rank stability was assessed separately by computing site ranks independently within each fold (Fig. S3, S7). Because correlated sequence features can share attribution, site ranks are used for prioritization rather than residue-level causal estimation. Full details of the SHAP aggregation procedure are in Supplementary Methods S1.6.

### 4.6 Cross-study comparison construction

The Koel set comprised the seven positions (145, 155, 156, 158, 159, 189, 193) identified as drivers of major H3N2 cluster transitions [11]. The Neher/Bedford reference set comprised 15 cluster-transition-associated positions [13]. The Harvey reference set comprised 15 high-confidence or top-25 posterior-inclusion positions from the Harvey et al. Bayesian analysis [14]. The Shah reference set comprised 30 positions identified by aggregating the top 20 feature-importance sites across 14 influenza seasons in the Shah et al. seasonal AdaBoost model trained on WIC HI data [15]. Additional sequence-only rankings included two Light-GBM+SHAP rankings generated from passage-history and collection-date prediction tasks without HI measurements [35], plus unpassaged sequence entropy and dN/dS rankings drawn from prior sequence analyses [17, 25]. For all comparisons, overlap was defined as the count of positions shared between the primary model’s top 30 and the reference set or external top-30 ranking. As a reference for interpreting overlap counts, the expected overlap under a hypergeometric null depends on the analysis universe: for the filtered WIC HA1 analysis, two independently drawn sets of 30 from 328 sites would overlap by 2.7 sites on average (0.6 for a 7-site benchmark; 1.4 for a 15-site benchmark), whereas for the Neher/Bedford mature-HA analysis the corresponding expectations over 550 sites are 1.6, 0.4, and 0.8. Full definitions of each reference set and external ranking are in Supplementary Methods S1.7. Benchmark overlap counts for all model variants are in Table S2; external-ranking overlaps are shown in Fig. 3B.

## Data Availability

Analysis configuration files, non-sequence input tables, manifests, cross-study comparison tables, and all manuscript figures are available in the public repository associated with this project. Raw HA sequence FASTA files are excluded from the public repository because redistribution is restricted by source-database terms, including those of GISAID. Non-sequence metadata sufficient to reproduce sequence retrieval is provided in the repository manifest files.

## Author Contributions

A.M. designed the study, performed the analyses, interpreted the results, and wrote the manuscript. M.S. interpreted the results and wrote the manuscript.

## Acknowledgments

We gratefully acknowledge the GISAID Initiative and all data contributors for making influenza sequence data publicly available. We also acknowledge the WHO Collaborating Centres for Reference and Research on Influenza and the laboratories that generated the HI data used in the WIC dataset.

## References

[1] Robert G. Webster, William J. Bean, Owen T. Gorman, Thomas M. Chambers, and Yoshihiro Kawaoka. Evolution and ecology of influenza A viruses. Microbiol. Rev., 56: 152–179, 1992.

[2] Nancy J. Cox and Kanta Subbarao. Global epidemiology of influenza: past and present. Annu. Rev. Med., 51:407–421, 2000.

[3] Derek J. Smith, Alan S. Lapedes, Jan C. de Jong, Theo M. Bestebroer, Guus F. Rimmelzwaan, Albert D. M. E. Osterhaus, and Ron A. M. Fouchier. Mapping the antigenic and genetic evolution of influenza virus. Science, 305:371–376, 2004.

[4] Colin A. Russell, Terry C. Jones, Ian G. Barr, Nancy J. Cox, Rebecca J. Garten, Vicky Gregory, Ian D. Gust, Alan W. Hampson, Alan J. Hay, Aeron C. Hurt, Jan C. de Jong, Anne Kelso, Alexander I. Klimov, Tsutomu Kageyama, Naomi Komadina, Alan S. Lapedes, Yi P. Lin, Ana Mosterin, Masatsugu Obuchi, Takato Odagiri, Albert D. M. E. Osterhaus, Guus F. Rimmelzwaan, Michael W. Shaw, Eugene Skepner, Klaus Stohr,Masato Tashiro, Ron A. M. Fouchier, and Derek J. Smith. The global circulation of seasonal influenza A(H3N2) viruses. Science, 320:340–346, 2008.

[5] Trevor Bedford, Steven Riley, Ian G. Barr, Shobha Broor, Mandeep Chadha, Nancy J. Cox, Rodney S. Daniels, C. Palani Gunasekaran, Aeron C. Hurt, Anne Kelso, Alexander Klimov, Nicola S. Lewis, Xiyan Li, John W. McCauley, Takato Odagiri, Varsha Potdar, Andrew Rambaut, Yuelong Shu, Eugene Skepner, Derek J. Smith, Marc A. Suchard, Masato Tashiro, Dayan Wang, Xiyan Xu, Philippe Lemey, and Colin A. Russell. Global circulation patterns of seasonal influenza viruses vary with antigenic drift. Nature, 523: 217–220, 2015.

[6] Robert G. Webster and W. Graeme Laver. Determination of the number of nonoverlapping antigenic areas on Hong Kong (H3N2) influenza virus hemagglutinin with monoclonal antibodies and the selection of variants with potential epidemiological significance. Virology, 104:139–148, 1980.

[7] Don C. Wiley, Ian A. Wilson, and John J. Skehel. Structural identification of the antibody-binding sites of Hong Kong influenza haemagglutinin and their involvement in antigenic variation. Nature, 289:373–378, 1981.

[8] Ian A. Wilson, John J. Skehel, and Don C. Wiley. Structure of the haemagglutinin membrane glycoprotein of influenza virus at 3 A resolution. Nature, 289:366–373, 1981.

[9] Walter Gerhard, Jonathan Yewdell, Marvin E. Frankel, and Robert Webster. Antigenic structure of influenza virus haemagglutinin defined by hybridoma antibodies. Nature, 290:713–717, 1981.

[10] Robin M. Bush, Catherine A. Bender, Kanta Subbarao, Nancy J. Cox, and Walter M. Fitch. Predicting the evolution of human influenza A. Science, 286:1921–1925, 1999.

[11] Björn F. Koel, David F. Burke, Theo M. Bestebroer, Stefan van der Vliet, GerbenC. M. Zondag, Gaby Vervaet, Eugene Skepner, Nicola S. Lewis, Monique I. J. Spronken,Colin A. Russell, Mikhail Y. Eropkin, Aeron C. Hurt, Ian G. Barr, Jan C. de Jong, Guus F. Rimmelzwaan, Albert D. M. E. Osterhaus, Ron A. M. Fouchier, and Derek J. Smith. Substitutions near the receptor binding site determine major antigenic change during influenza virus evolution. Science, 342:976–979, 2013.

[12] Trevor Bedford, Marc A. Suchard, Philippe Lemey, Gytis Dudas, Victoria Gregory, Alan J. Hay, John W. McCauley, Colin A. Russell, Derek J. Smith, and Andrew Rambaut. Integrating influenza antigenic dynamics with molecular evolution. eLife, 3: e01914, 2014.

[13] Richard A. Neher, Trevor Bedford, Rodney S. Daniels, Colin A. Russell, and Boris I. Shraiman. Prediction, dynamics, and visualization of antigenic phenotypes of seasonal influenza viruses. Proc. Natl. Acad. Sci. USA, 113:E1701–E1709, 2016.

[14] William T. Harvey, Vinny Davies, Rodney S. Daniels, Lynne Whittaker, Victoria Gregory, Alan J. Hay, Dirk Husmeier, John W. McCauley, and Richard Reeve. A Bayesian approach to incorporate structural data into the mapping of genotype to antigenic phenotype of influenza A(H3N2) viruses. PLoS Comput. Biol., 19:e1010885, 2023.

[15] Syed Awais W. Shah, Daniel P. Palomar, Ian Barr, Leo L. M. Poon, Ahmed Abdul Quadeer, and Matthew R. McKay. Seasonal antigenic prediction of influenza A H3N2 using machine learning. Nat. Commun., 15:3833, 2024.

[16] Robin M. Bush, Catherine B. Smith, Nancy J. Cox, and Walter M. Fitch. Effects of passage history and sampling bias on phylogenetic reconstruction of human influenza A evolution. Proc. Natl. Acad. Sci. USA, 97:6974–6980, 2000.

[17] Claire D. McWhite, Austin G. Meyer, and Claus O. Wilke. Sequence amplification via cell passaging creates spurious signals of positive adaptation in influenza virus H3N2 hemagglutinin. Virus Evol., 2:vew026, 2016.

[18] Lauren Parker, Stephen A. Wharton, Stephen R. Martin, Katie Cross, Yipu Lin, Yan Liu, Ten Feizi, Rodney S. Daniels, and John W. McCauley. Effects of egg-adaptation on receptor-binding and antigenic properties of recent influenza A (H3N2) vaccine viruses. J. Gen. Virol., 97:1333–1344, 2016.

[19] Seth J. Zost, Kaela Parkhouse, Megan E. Gumina, Kangchon Kim, Sebastian Diaz Perez, Patrick C. Wilson, John J. Treanor, Andrea J. Sant, Sarah Cobey, and Scott E. Hensley. Contemporary H3N2 influenza viruses have a glycosylation site that alters binding of antibodies elicited by egg-adapted vaccine strains. Proc. Natl. Acad. Sci. USA, 114:12578–12583, 2017.

[20] Kathryn Kistler and Trevor Bedford. Seasonal influenza viruses show distinct adaptive dynamics during growth in chicken eggs. Mol. Biol. Evol., 42:msaf227, 2025.

[21] Walter M. Fitch, Jodi M. E. Leiter, Xianqun Li, and Peter Palese. Positive Darwinian evolution in human influenza A viruses. Proc. Natl. Acad. Sci. USA, 88:4270–4274, 1991.

[22] Masatoshi Nei and Takashi Gojobori. Simple methods for estimating the numbers of synonymous and nonsynonymous nucleotide substitutions. Mol. Biol. Evol., 3:418–426, 1986.

[23] Yoshiyuki Suzuki. Natural selection on the influenza virus genome. Mol. Biol. Evol., 23:1902–1911, 2006.

[24] Keyao Pan and Michael W. Deem. Quantifying selection and diversity in viruses by entropy methods, with application to the haemagglutinin of H3N2 influenza. J. R. Soc. Interface, 8:1644–1653, 2011.

[25] Austin G. Meyer and Claus O. Wilke. Geometric constraints dominate the antigenic evolution of influenza H3N2 hemagglutinin. PLOS Pathog., 11:e1004940, 2015.

[26] Julian Echave, Stephanie J. Spielman, and Claus O. Wilke. Causes of evolutionary rate variation among protein sites. Nat. Rev. Genet., 17:109–121, 2016.

[27] Austin G. Meyer, Stephanie J. Spielman, Trevor Bedford, and Claus O. Wilke. Time dependence of evolutionary metrics during the 2009 pandemic influenza virus outbreak. Virus Evol., 1:vev006, 2015.

[28] Simon Y. W. Ho, Matthew J. Phillips, Alan Cooper, and Alexei J. Drummond. Time dependency of molecular rate estimates and systematic overestimation of recent divergence times. Mol. Biol. Evol., 22:1561–1568, 2005.

[29] Yuan-Ling Xia, Weihua Li, Yongping Li, Xing-Lai Ji, Yun-Xin Fu, and Shu-Qun Liu. A deep learning approach for predicting antigenic variation of influenza A H3N2. Comput. Math. Methods Med., 2021:9997669, 2021.

[30] Marta Luksza and Michael Lässig. A predictive fitness model for influenza. Nature, 507: 57–61, 2014.

[31] Lars Steinbrück, Thorsten R. Klingen, and Alice C. McHardy. Computational prediction of vaccine strains for human influenza A (H3N2) viruses. J. Virol., 88:12123–12132, 2014.

[32] Brian Hie, Ellen D. Zhong, Bonnie Berger, and Bryan Bryson. Learning the language of viral evolution and escape. Science, 371:284–288, 2021.

[33] Nicole N. Thadani, Sarah Gurev, Pascal Notin, Noor Youssef, Nathan J. Rollins, Daniel Ritter, Chris Sander, Yarin Gal, and Debora S. Marks. Learning from prepandemic data to forecast viral escape. Nature, 622:818–825, 2023.

[34] Brent E. Allman, Luiz Vieira, Daniel J. Diaz, and Claus O. Wilke. A systematic evaluation of the language-of-viral-escape model using multiple machine learning frameworks. J. R. Soc. Interface, 22:20240598, 2025.

[35] Austin G. Meyer. Explainable machine learning reveals evolutionary signals in Influenza hemagglutinin. In Press at J. R. Soc. Interface, bioRxiv preprint available, 2026. doi: 10.1101/2025.09.21.677610.

[36] BjÖrn F. Koel, David F. Burke, Theo M. Bestebroer, Stefan van der Vliet, Gerben C. M. Zondag, Gaby Vervaet, Eugene Skepner, Nicola S. Lewis, Monique I. J. Spronken, Colin A. Russell, Mikhail Y. Eropkin, Aeron C. Hurt, Ian G. Barr, Jan C. de Jong, Guus F. Rimmelzwaan, Albert D. M. E. Osterhaus, Ron A. M. Fouchier, and Derek J. Smith. Epistatic interactions can moderate the antigenic effect of substitutions in haemagglutinin of influenza H3N2 virus. J. Gen. Virol., 100:773–777, 2019.

[37] Sergey Kryazhimskiy, Jonathan Dushoff, Georgii A. Bazykin, and Joshua B. Plotkin. Prevalence of epistasis in the evolution of influenza A surface proteins. PLoS Genet., 7: e1001301, 2011.

[38] Benjamin S. Chambers, Kaela Parkhouse, Ted M. Ross, Kevin Alby, and Scott E. Hensley. Identification of hemagglutinin residues responsible for H3N2 antigenic drift during the 2014–2015 influenza season. Cell Rep., 12:1–6, 2015.

[39] Austin G. Meyer and Claus O. Wilke. Integrating sequence variation and protein structure to identify sites under selection. Mol. Biol. Evol., 30:36–44, 2013.

[40] Suman R. Das, Scott E. Hensley, Alexandre David, Leah Schmidt, James S. Gibbs, Kanta Puig, Jack R. Bennink, and Jonathan W. Yewdell. Fitness costs limit influenza A virus hemagglutinin glycosylation as an immune evasion strategy. Proc. Natl. Acad. Sci. USA, 108:E1417–E1422, 2011.

[41] Ivan Kosik, William L. Ince, Lauren E. Gentles, Andrew J. Oler, Martina Kosikova, Matthew Angel, Javier G. Magadán, Pranav Bhatt, Nicole Wohlgemuth, Benjamin J. Cowling, Hana Golding, Surender Khurana, and Jonathan W. Yewdell. Influenza A virus hemagglutinin glycosylation compensates for antibody escape fitness costs. PLoS Pathog., 14:e1006796, 2018.

[42] Meghan O. Altman, Matthew Angel, Ivan Kosik, Nivaldo S. Trovao, Sun-Woo Yoon, Hana Golding, William L. Ince, Ian Crozier, Ruben O. Donis, Scott E. Hensley, Matthew J. Memoli, and Jonathan W. Yewdell. Human influenza A virus hemagglutinin glycan evolution follows a temporal pattern to a glycan limit. mBio, 10:e00204–19, 2019.

[43] Juhye M. Lee, John Huddleston, Michael B. Doud, Kathryn A. Hooper, Nicholas C. Wu, Trevor Bedford, and Jesse D. Bloom. Deep mutational scanning of hemagglutinin helps predict evolutionary fates of human H3N2 influenza variants. Proc. Natl. Acad. Sci. USA, 115:E8276–E8285, 2018.

[44] Bargavi Thyagarajan and Jesse D. Bloom. The inherent mutational tolerance and antigenic evolvability of influenza hemagglutinin. eLife, 3:e03300, 2014.

[45] Stefan Elbe and Gemma Buckland-Merrett. Data, disease and diplomacy: GISAID’s innovative contribution to global health. Glob. Chall., 1:33–46, 2017.

[46] Yuelong Shu and John McCauley. GISAID: Global initiative on sharing all influenza data – from vision to reality. Euro Surveill., 22:30494, 2017.

[47] Kazutaka Katoh and Daron M. Standley. MAFFT multiple sequence alignment software version 7: improvements in performance and usability. Mol. Biol. Evol., 30:772–780, 2013.

[48] Guolin Ke, Qi Meng, Thomas Finley, Taifeng Wang, Wei Chen, Weidong Ma, Qiwei Ye, and Tie-Yan Liu. LightGBM: A highly efficient gradient boosting decision tree. Adv. Neural Inf. Process. Syst., 30:3146–3154, 2017.

[49] Morgan N. Price, Paramvir S. Dehal, and Adam P. Arkin. FastTree 2–approximately maximum-likelihood trees for large alignments. PLoS One, 5:e9490, 2010.

[50] Peter J. A. Cock, Tiago Antao, Jeffrey T. Chang, Brad A. Chapman, Cymon J. Cox, Andrew Dalke, Iddo Friedberg, Thomas Hamelryck, Frank Kauff, Bartek Wilczynski, and Michiel J. L. de Hoon. Biopython: freely available Python tools for computational molecular biology and bioinformatics. Bioinformatics, 25:1422–1423, 2009.

[51] Scott M. Lundberg and Su-In Lee. A unified approach to interpreting model predictions. Adv. Neural Inf. Process. Syst., 30:4765–4774, 2017.

